# Subclonal evolution and expansion of spatially distinct THY1-positive cells is associated with recurrence in glioblastoma

**DOI:** 10.1101/2021.09.10.459454

**Authors:** Wajd N. Al-Holou, Hanxiao Wang, Visweswaran Ravikumar, Morgan Oneka, Roel GW Verhaak, Hoon Kim, Drew Pratt, Sandra Camelo-Piragua, Corey Speers, Daniel R Wahl, Sunita Shankar, Todd Hollon, Oren Sagher, Jason A Heth, Karin M. Muraszko, Theodore S. Lawrence, Ana C de Carvalho, Tom Mikkelsen, Arvind Rao, Alnawaz Rehemtulla

## Abstract

**Purpose:** Glioblastoma (GBM) is a lethal disease characterized by inevitable recurrence. Here we investigate the molecular pathways mediating resistance, with the goal of identifying therapeutic opportunities to target this tumor.

**Experimental Design:** We developed a longitudinal *in vivo* recurrence model utilizing patient-derived explants to produce paired specimens (pre- and post-recurrence) following temozolomide(TMZ) and radiation(IR). These specimens were evaluated for treatment response and to identify gene expression pathways driving treatment resistance. Findings were clinically validated using spatial transcriptomics of human GBMs.

**Results:** These studies reveal in replicate cohorts, a gene expression profile characterized by upregulation of mesenchymal and stem-like genes at recurrence. Analyses of clinical databases revealed increased expression of this transcriptional profile to be significantly associated with worse median overall survival (248 days vs 430 days, p=0.0004), and upregulation of this profile at recurrence. Most notably, we identified upregulation of *TGFβ* signaling, and more than one-hundred-fold increase in *THY1* levels at recurrence. Utilizing cell sorting, we observed that THY1-positive cells represented <10% of cells in the treatment-naïve tumors and 75-96% in the recurrent tumors. We then isolated THY1-positive cells from treatment-naïve patient samples and determined that they were inherently resistant to chemoradiation in orthotopic models. Additionally, using image-guided biopsies from treatment-naïve human GBM, we conducted spatial transcriptomic analyses. This revealed rare *THY1*+ regions characterized by mesenchymal/stem-like gene expression, analogous to our recurrent mouse model samples, which co-localized with macrophages within the perivascular niche. Since TGFβ signaling contributes to a mesenchymal/stem-like phenotype, we inhibited TGFβRI activity *in vivo* which resulted in decreased mesenchymal/stem-like protein levels, including THY1, and restored sensitivity to TMZ/IR in recurrent tumors.

**Conclusions:** These findings reveal that GBM recurrence may result from tumor repopulation by pre-existing, therapy-resistant, THY1-positive, mesenchymal/stem-like cells within the perivascular niche. Furthermore, our data demonstrate the promise of targeting upregulated pathways in resistant subclones as a novel mechanism to achieve therapeutic response, and specifically that THY1 expression may represent a biomarker of response to TGFβ inhibition.

## INTRODUCTION

Glioblastoma (GBM), the most common primary intraparenchymal brain tumor, is a lethal disease with a median survival of 15 months that is characterized by treatment resistance, aggressive brain invasion, and inevitable recurrence, with less than 10% of patients surviving beyond 5 years[1, 2]. A major obstacle contributing to rapid recurrence and the lack of effective treatments is the marked intra- and inter-tumoral heterogeneity in GBM[3–6]. Furthermore, given the heterogeneity present within GBM, it has been postulated that evolution and expansion of pre-existing treatment-resistant sub-clones are often responsible for inevitable recurrence [5, 7, 8]. Intratumoral heterogeneity can be seen at both at the cellular and microenvironmental level. At the tumor microenvironmental level, distinct niches, especially perivascular, hypoxic, and invasive tumor niches, have been shown to be enriched with therapy-resistant cancer stem cell (CSC) populations, also described as recurrence-initiating stem cells [3, 7, 9, 10]. These pre-existing CSC populations are thought to play a critical role in driving treatment resistance and are characterized by self-renewal, differentiation, and the ability to expand and repopulate tumors post-therapy[11]. Definitive evidence for the role for pre-existing treatment-resistant cancer stem cells in evading chemo-radiation therapy as well as the mechanistic basis for this phenotype is lacking. An understanding of the genetic basis for the therapy resistant phenotype should provide an opportunity for therapeutic intervention to decrease the recurrence rate in GBM.

In this effort, we have utilized a longitudinal mouse model developed using patient-derived GBM explants to recapitulate the clinical treatment paradigm. Using this model, we generated pre-treatment and recurrent paired samples from each experimental mouse with the goal of identifying differentially expressed genes that should provide a molecular basis for tumor recurrence. Systematic analyses of multiple replicate intracranial patient-derived xenografts(PDX) revealed that recurrent tumors were characterized by a gene expression pattern characteristic of a mesenchymal and CSC signature, which include upregulation of *THY1/CD90, TGFβ1, TGFβ2, SOX2, ZEB2, and GLI2*. Consistent with previous observations, we demonstrate using a large GBM cohort that this gene expression profile is associated with a worse overall prognosis, highlighting the clinical implications of our findings. To test the hypothesis that tumor recurrence may be attributed to a pre-existing cell population characterized by this gene expression signature, we immuno-sorted THY1+ cells from treatment-naïve patient samples and demonstrated that intracranial tumors derived from THY1+ cells (Pre-THY1+) were inherently treatment resistant compared to unsorted tumors. These results suggest that rare populations of inherently treatment-resistant cells with distinct gene expression profiles can be found within the original treatment-naïve tumors, and are responsible for tumor recurrence[5, 8, 12]. In support, treatment of tumors with a targeted agent that reverses the mesenchymal gene expression signature, resulted in restoration of sensitivity to chemoradiation. Furthermore, spatial transcriptomic analyses of human GBM specimens consistently identified *THY1*+ cells co-localized with expression of mesenchymal, glioma stem cell, and macrophage gene signatures and were primarily found within a perivascular niche. This suggests an important role for these rare resistant cells in the vascular tumor micro-environment.

## MATERIALS AND METHODS

### Patient-derived primary neurosphere culture

Low passage, patient-derived primary GBM neurosphere cells, HF2303, have been previously described[13, 14], and were maintained in neurosphere medium composed of DMEM/F12 medium (#11320-033, Life Technologies, Carlsbad, CA) plus N2 supplement(#17502-048, Life Technologies, Carlsbad, CA), 0.5mg/ml BSA (#A4919, Sigma-Aldrich Co., St. Louis, MO), 25µg/ml Gentamicin (#15750-060, Life Technologies, Carlsbad, CA), 0.5% antibiotic/antimycotic (#15240-096, Life Technologies, Carlsbad, CA), 20ng/ml bFGF and 20 ng/ml EGF (#100-15 and #100-18B, PeproTech, Rocky Hill NJ). Cultures were derived from resected brain tumor specimens collected at Henry Ford Hospital (Detroit, MI) with written informed consent from patients, under a protocol approved by the Institutional Review Board.

### Stereotactic biopsy

Coordinates for MRI-guided biopsy were determined by a fiducial markers attached to the whole-body volume transmit coil and VnmrJ software (Agilent Technologies, Inc., Santa Clara, CA)[15] (Fig. S1). The mouse was moved to a stereotactic station along with the whole-body volume transmit coil. A 1cm incision was made and the skull was exposed with cotton-tip applicators. After biopsy location was recovered using the fiducial markers, a burr hole was drilled and a 22GA x 3 7/8” needle (#54722, Inrad, Kentwood, MI) attached to a vacuum syringe was inserted into the tumor. Biopsy tissues were dissociated and cultured in NMGF medium immediately and were expanded for three passages, to eliminate non-dividing cells derived from normal mouse brain. Holes were covered using bonewax and incisions were sealed using Vetbond. 100ul of carprofen was subcutaneously injected following the surgical procedure.

### Treatment

For mice bearing intracranial tumors, animals were randomized into treatment groups once their tumor volume reached 20-30mm^3^ by MRI evaluation(Fig. 1). Temozolomide (LKT Laboratories, St Paul, MN) was suspended in Ora-Plus suspending vehicle (#0574-0303-16, Rotterdam, Netherlands) and administered to animals (66mg/kg) via oral gavage daily for five days per week. Cranial irradiation was carried out one hour post temozolomide treatment. Mice were restrained in a home-made plastic restraining device. A lead shield was used so that only the head was exposed to radiation using a Kimtron INC-320 orthovoltage irradiator (Kimtron Medical, Oxford, CT). A total of 20Gy radiation was delivered to each animal at 2 Gy/day for ten days. Maintenance temozolomide was delivered orally three times week every other week from the third week till animals became moribund. For mice with subcutaneous flank tumors, animals were randomized into study groups when subcutaneous tumors reached an approximate volume of 300 mm^3^. Temozolomide and radiation were delivered via similar administration routes and treatment schedules as described above. Radiation was targeted at local tumor sites with a lead shield. LY2109761(50mg/kg) (AbMole, Houston, TX), a TGFβRI inhibitor, was reconstituted using Ora-Plus suspending vehicle and delivered to animals twice per day for five days every week for two weeks. For animals treated with temozolomide, LY2109761 and radiation, LY2109761 was administered one hour before radiation along with TMZ and six hours after radiation treatment.

**Fig. 1.**
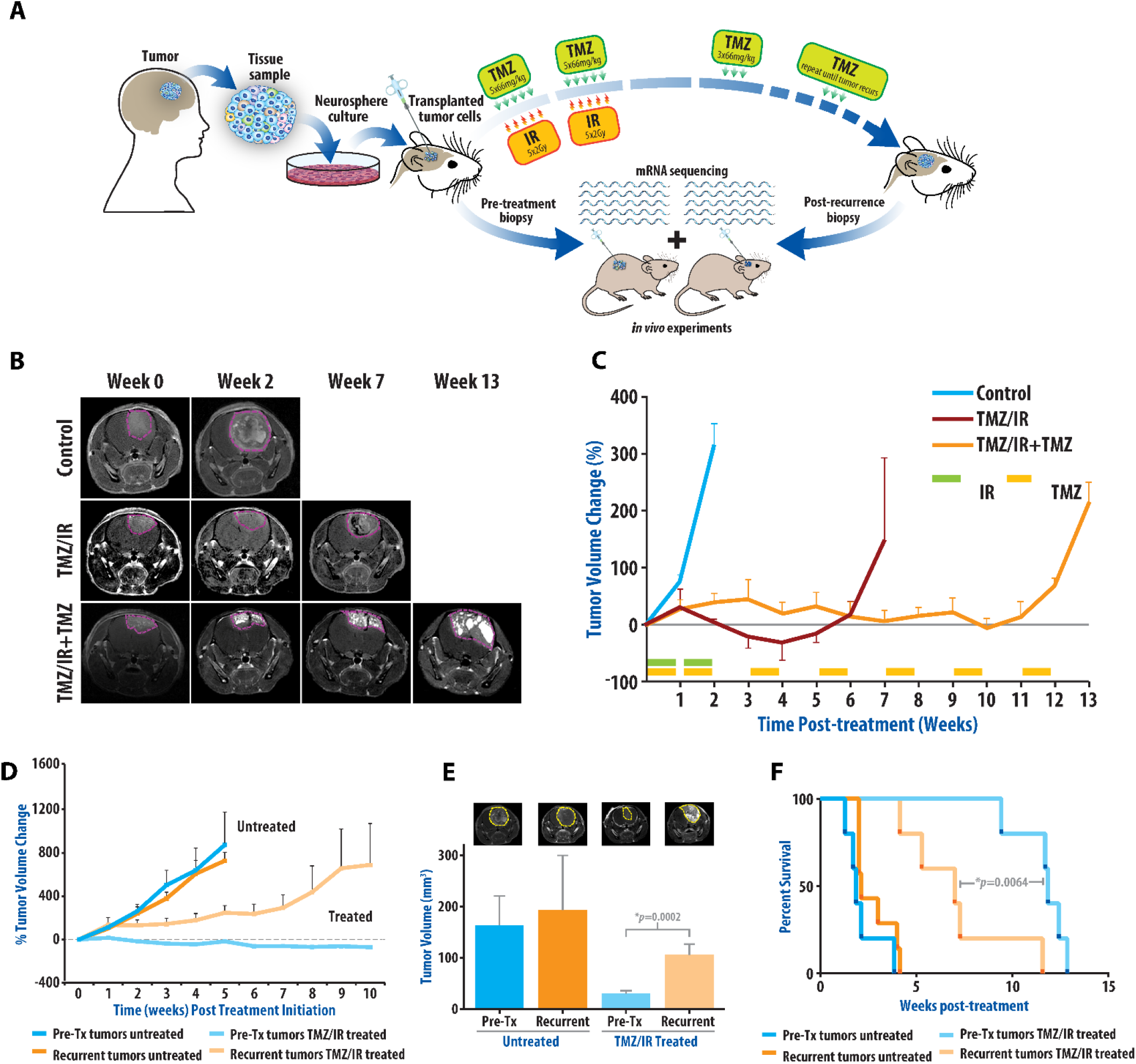
Analysis of longitudinal intracranial GBM samples, pre-treatment and at recurrence, reveals development of a therapeutic resistant phenotype. **(A)** Schematic of experimental design. Mice bearing intracranial tumors derived from human GBM explants are treated with concomitant temozolomide (TMZ) and radiation (IR) for two weeks followed by adjuvant TMZ until animals became moribund. Tumor biopsies are obtained pre-treatment and at recurrence to provide samples for molecular analysis and additional *in vivo* and *in vitro* comparative studies. **(B)** Representative MR images from Control, TMZ/IR and TMZ/IR+TMZ treatment groups at 0, 2, 7 and 13 weeks post-initiation. **(C)** Treatment response quantified using MR-imaging derived tumor volume changes in Control, TMZ/IR and TMZ/IR+TMZ intracranial tumor bearing groups. TMZ/IR treatment involved 2 weeks of concomitant TMZ/IR, while mice receiving TMZ/IR+TMZ received the initial 2 week treatment followed by maintenance adjuvant TMZ every other week. Percent change in tumor volume is shown as mean change in a cohort of five animals with SEM. **(D)** Pre-treatment(Pre-Tx) and post-treatment(recurrent) biopsy samples were re-implanted subcutaneously into the flanks of mice to evaluate their sensitivity to treatment as described in methods. Pre-treatment and recurrent tumors show similar growth rates in the absence of treatment, while recurrent tumors exhibit an inherent resistance to TMZ/IR. **(E)** Samples described in D were also evaluated intracranially. Tumor volumes derived from pre-treatment and recurrent tumors with or without TMZ/IR are shown as mean tumor volumes (+SEM) as well as representative MR images three weeks after treatment initiation. **(F)** Kaplan-Meier survival curve of mice after intracranial implantation of pre-treatment and recurrent tumors with or without TMZ/IR treatment. Mice bearing recurrent tumors have significantly worse survival with treatment.

### Spatial transcriptomics

Under an institutional review board approved study, intraoperative brain tumor specimens were collected at the University of Michigan using MRI-imaged guidance technology to localize specimen within contrast-enhanced tissue with low and high perfusion parameters. This research was in full compliance of all pertinent ethical regulations for research with human biospecimens and all data were de-identified (IRB HUM00175135). Human samples acquired were immediately frozen in liquid nitrogen/isopentane, after which they were imbedded in optimal cutting temperature compound and frozen. Tissue was sectioned onto Visium Spatial Gene Expression Slide(10X Genomics) and permeabilized to release mRNA to bind to spatially barcoded capture probes. This was used to synthesize cDNA followed by preparation of sequencing libraries and sequencing. Sequencing data was visualized using and the resulting images put together with Vslide (v1.1.115, MetaSystems GmbH). Sample data are imaged and analyzed with 10X genomics software(Space Ranger and Loupe Browser). The SpaceRanger output files were imported into R using the Load10x_Spatial function. Library size normalization for the spots was performed using the SCTransform function[16], controlling for the percent of mitochondrial genes expressed at each spot. We imputed gene counts using MAGIC to smooth out the data prior to performing dimension reduction using PCA and UMAP[17]. Specifically, the principal components were computed using top variable genes after imputation, and embedding of data on the first 30 principal components is used for identifying clusters and learning UMAP embedding. Pathway enrichment scores for specific genesets are computed similar to Neftel *et al*[18]. A geneset collection object was prepared by collating specific genesets of interest from MSigDb as well as other datasets[19, 20]. Enrichment scores were computed based on the MAGIC-imputed counts data. All genes were grouped into 30 bins based on mean expression. Within each given gene set, mean expression of each gene was computed along with one hundred other randomly sampled genes from the corresponding bins as target genes. The difference in mean expression of target genes of interest versus control genes was calculated at each spot to compute the enrichment scores. After enrichment scores for the genesets at each spot are calculated, we then generated scatterplots of pathway scores versus *THY1* expression, and computed Spearman partial correlations between the two controlling for the G1.S cell cycle score at the spots, to account for variations in cell viability across the tissue section. To determine spatial overlap between *THY1* expression and pathway activity, we performed a regional Fisher’s exact test for each spot. Based on *THY1* expression and pathway scores, we binarized the data into THY1-positive and pathway-positive at their 75th quantile. We then reviewed the k=25 nearest neighbors for every spot, and performed a Fisher’s exact test to determine if there is significant overlap of the two features, i.e. if the pathway scores happen to be specifically enriched at THY1-positive regions. The Fisher test p-values are then overlaid onto each spot on the tissue slice to visualize regions that have significant overlaps in the two features.

### Statistics

Results of *in vivo* volume data are shown as mean with standard error with 5 replicates performed for studies. Student’s t-test (unpaired) and ANOVA was performed to assess statistical significance with p<0.05 considered significant. Survival analyses were performed utilizing Kaplan-Meier survival and statistical significance tested utilizing log-rank (Mantel-Cox) analyses with p-values as shown utilizing Graphpad Prism(9.0). Assessment of publicly accessible data from The Cancer Genome Atlas and Glass Consortium[21] was analyzed for gene sets of interest.

## RESULTS

### *in vivo* modeling recapitulates therapeutic response to TMZ/IR followed by recurrence

To evaluate the therapeutic response and the underlying genetic basis for resistance to standard of care therapy (TMZ/IR) in GBM, we utilized patient derived explants in intracranial xenografts (Fig. 1). Magnetic resonance imaging was performed weekly to monitor tumor growth after implantation. Mice bearing tumors of approximately 20mm^3^ were randomized into three study cohorts. The first represented control untreated animals, and a second cohort was treated with concurrent TMZ/IR for two weeks. A third cohort was administered concurrent TMZ/IR for 2 weeks followed by adjuvant TMZ three times a week, every other week after completion of the second week of TMZ/IR. This experimental design was intended to investigate mechanisms of resistance to standard of care therapy (Fig. 1A). In contrast to control animals which exhibited a greater than 300% increase in tumor volume and succumbed to disease by the third week, TMZ/IR treated animals exhibited an initial tumor regression and survived beyond six weeks. Mice that received adjuvant TMZ exhibited a greater delay in tumor growth and had a prolonged survival (>12 weeks). Despite an initial response to therapy, all treated animals had recurrence (Fig. 1B and 1C).

To better understand the underlying genetic basis for the development of therapeutic resistance, an MRI-guided stereotactic intracranial biopsy was used to obtain tumor tissue from animals prior to initiation of treatment. This biopsy procedure does not significantly impact the gross tumor architecture[15](Fig. S1). Viable tumor tissue was also obtained from these animals upon recurrence. The paired pre-treatment and recurrent tumor samples were collected for further analyses.

### Recurrent tumors are distinctly resistant to TMZ/IR

To evaluate the sensitivity of recurrent tumors to therapy, paired neurosphere cultures derived from pre-treatment and recurrent biopsies were implanted intracranially and subcutaneously into immune-deficient mice. Subcutaneously implanted murine models with palpable tumors were randomized into a control group or treated with TMZ/IR for two weeks followed by adjuvant TMZ as above. In the absence of treatment, both the pre-treatment and recurrent tumors obtained by biopsies grew at a similar rate. In response to TMZ/IR, pre-treatment tumors demonstrated a significant delay in tumor growth (analogous to the initial patient-derived tumors), and exhibited a 50% decrease in tumor volume. However, tumors derived from recurrent biopsies continued to exhibit resistance to TMZ/IR, and demonstrated a 750% increase in tumor volume (Fig. 1D). To confirm these findings in an appropriate tumor microenvironment, intracranial tumors were established using the same paired neurospheres. Treatment of mice bearing intracranial recurrent tumors with TMZ/IR plus adjuvant TMZ demonstrated their resistance to therapy compared to those derived from pre-treatment samples (Fig. 1E). In support, the median survival of mice bearing recurrent tumors was 7 weeks compared to 11.9 weeks for treatment naïve tumors (p=0.006) (Fig. 1F).

### Recurrent tumors develop a therapeutic resistant phenotype with increased mesenchymal and stem cell gene expression patterns and *THY1* upregulation

To delineate the underlying genomic basis for the observed resistance to therapy in recurrent samples, transcriptomal analysis of at least three independent replicates of pre-treatment and recurrent biopsies from the same subject animals was performed using RNAseq(Fig. 2A). 1159 genes were found to be differentially expressed (FDR<0.05). Among these differentially expressed genes, 645 genes were upregulated in recurrent tumor samples while 514 genes were downregulated. Unsupervised clustering analysis showed distinct transcriptomic profiles of the pretreated and recurrent tumors across multiple replicates (Fig. 2B, Fig. S2). Moreover, the transcriptome of each pre-treatment sample closely resembled that of the original patient derived neurosphere cultures, demonstrating that gene expression profile did not drift significantly after *in vitro* expansion or upon intracranial implantation.

**Fig. 2.**
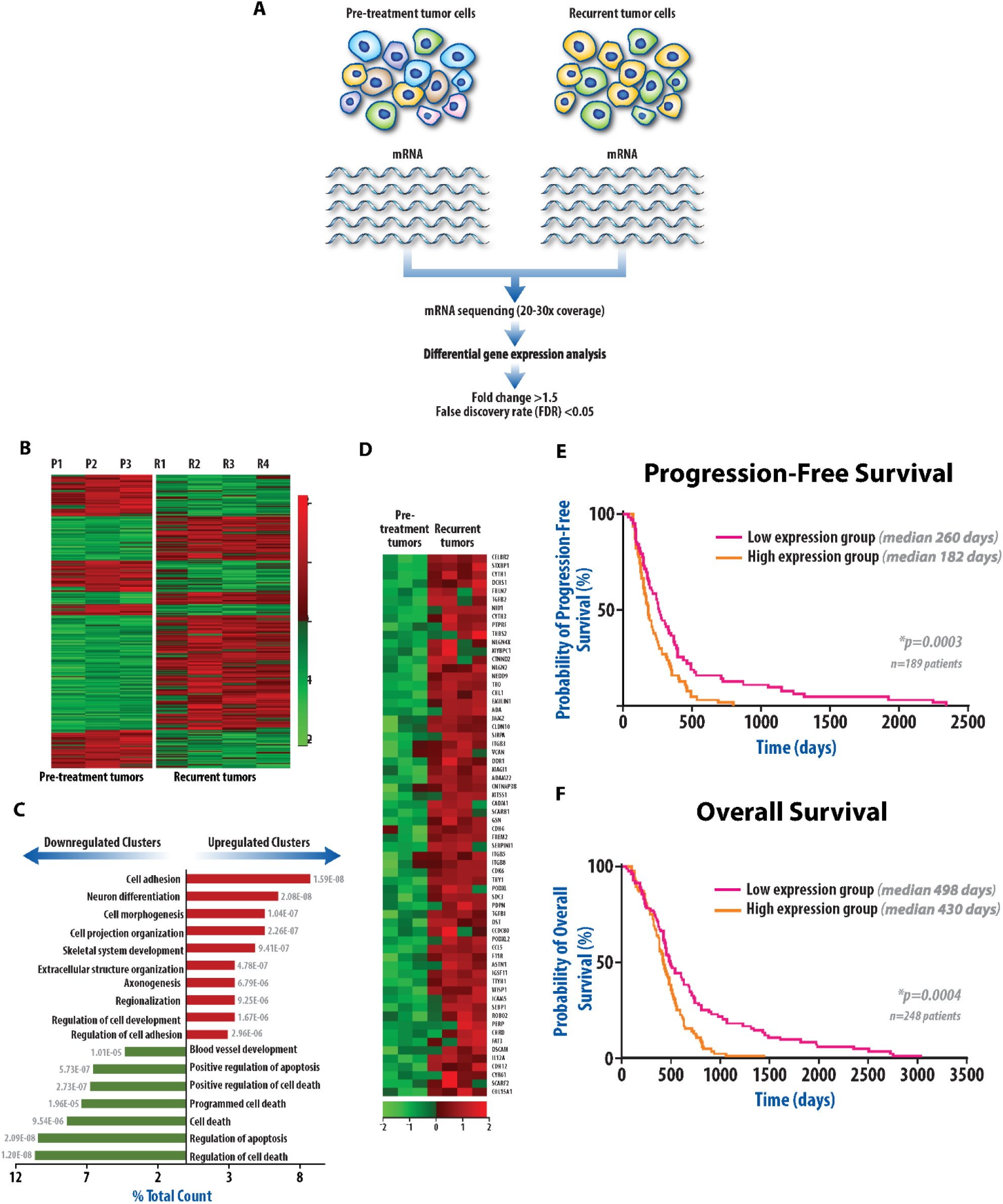
RNAseq analyses reveal a resistant gene expression profile that predicts outcomes in clinical datasets. **(A)** A diagrammatic representation of the experimental workflow. Paired pre-treatment and recurrent tumor samples are evaluated by next generation sequencing of RNA samples extracted from cells. **(B)** Unsupervised clustering analysis revealed that replicate independent pre-treatment samples clustered together had a similar gene expression pattern. Similarly independent replicate recurrent samples clustered together. Heatmap of differentially expressed genes in independent pre-treatment biopsies (P1,P2,P3) and recurrent tumor biopsies (R1,R2,R3,R4). **(C)** Functional clusters of upregulated and downregulated genes in recurrent tumors. Bar graph shows percentage of gene counts of each cluster compared to total upregulated or downregulated genes. P-value of each cluster is labeled next to each bar. Cell adhesion genes were upregulated in recurrent tumors. **(D)** Heatmap of individual genes within the highest upregulated clusters. Progression-free **(E)** and overall survival **(F)** and progression-free survival of GBM patients with high and low expression levels of a signature characterized by cell adhesion genes, showing that this upregulated signature predicts outcomes in GBM patients.

Next, we carried out a Gene Ontology analysis using Database for Annotation, Visualization and Integrated Discovery (DAVID) to identify cellular processes enriched in tumors with TMZ/IR resistance. Recurrent tumors showed altered biological processes as 20 upregulated and 13 downregulated functional clusters (FDR<0.05). These clusters were further consolidated to yield a non-redundant set of 10 upregulated gene clusters and seven downregulated gene clusters (Fig. 2C). Of the seven downregulated clusters, six were associated with apoptosis and cell death. Genes involved in cell adhesion, neuronal differentiation/development, and cellular morphogenesis were dominant in the upregulated clusters. Particularly, recurrent tumors overexpressed genes related to cell adhesion (Fig. 2D) (e.g. *ITGB3, ITGB5, ITGB8, NEDD9, FBLN7, NLGN2, CADM1 and VCAN*) and those associated with a mesenchymal phenotype (*TGFΒ2, TGFΒ1, THY1*).

Upregulation of this gene expression signature was found to be associated with a therapy resistant phenotype as indicated by a worse prognosis in a clinical database. A metagene score was created based on the average gene expression level of the top 64 cell adhesion genes. We selected GBM patients treated with TMZ and IR from the TCGA database and ranked them based on the metagene scores. Progression-free and overall survival were compared in the patients with a high metagene score (upper 1/3 total selected patients) and those with a low metagene score (lower 1/3 of total selected patients). As shown in Figs. 2E and 2F, patients with a high metagene score had worse clinical outcome, both in terms of median progression free survival (260 days compared to 189 days, p=0.0003), as well as median overall survival (median survival of 430 days compared to 248 days, p=0.0004). This analysis corroborates our observations from the experimental mouse model that an increase in the experimentally observed signature correlates with therapeutic resistance in patients.

To explore the role of genes associated with cell adhesion, mesenchymal and stem cell phenotype in TMZ/IR resistance, we first validated select genes from RNAseq analysis (*ZEB2, VCAN, CDK6, THY1, GLI2, SOX2*) which have previously been associated with therapeutic resistance in GBM[22–25]. Quantitative PCR analysis confirmed that *ZEB2, VCAN, CDK6, THY1, GLI2,* and *SOX2* mRNA levels were upregulated in all four recurrent tumor samples compared to their treatment naïve counterparts (Figs. 3A,B). Western blot analysis confirmed that the protein levels for each of these genes were also increased in a majority of recurrent samples compared to their untreated counterparts (Figs. 3C,D). Expression of the CD133 cell surface epitope, a marker of glioma stemness, was also markedly elevated in three of the four recurrent tumors (Fig. 3B).

**Fig. 3.**
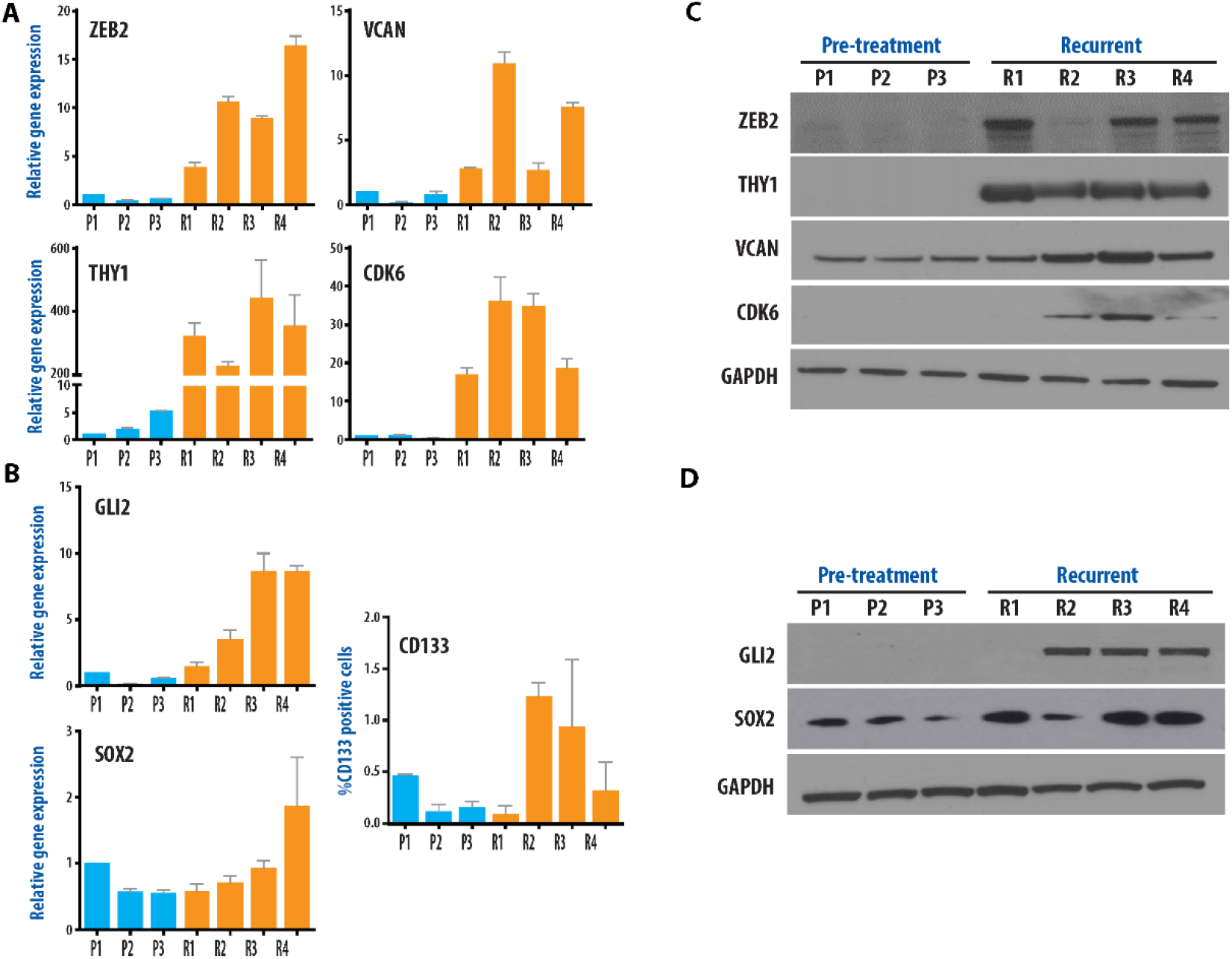
Analyses of paired samples reveals development of a therapeutic resistant phenotype with a mesenchymal, and stem cell gene expression pattern. **(A)** Upregulated genes in recurrent samples (R1-R4) associated with a mesenchymal signature were confirmed by qRTPCR, show notable upregulation in recurrent samples compared to pre-treatment samples (P1-P3). THY1 is most prominently differentially upregulated, greater than 100-fold compared to pre-treatment levels. **(B)** Upregulated genes in recurrent samples associated with a stem cell phenotype were confirmed using qRTPCR. FACS analysis of CD133 expression in pre-treatment and recurrent neurosphere cells (bottom right panel) shows upregulation of CD133+ cells in 3 of 4 samples at recurrence. **(C, D)** Western blot analysis confirmed that the protein levels for each of these genes were also increased in a majority of recurrent samples compared to their untreated counterparts.

In contrast to these consistent changes in gene expression observed between replicate recurrent tumors, analysis of copy number gain or copy number loss between each of the pre-treatment and recurrent samples did not show any significant and consistent changes (Fig. S3). This suggests that development of the resistance phenotype resulted from changes in gene expression rather than genomic changes, highlighting the importance of modulation of the relevant cell signaling pathways identified.

### Inhibition of TGFβ signaling decreases *THY1* expression, reverses the mesenchymal resistance phenotype, and results in improved treatment sensitivity *in vivo*

In our recurrent tumors, the resistance phenotype is characterized by increased expression of mesenchymal and CSC genes. Specifically, the TGFβ pathway was also significantly upregulated. In fact, many (21 of 63) of the upregulated genes within the recurrent tumors are associated directly or indirectly with the TGFβ signaling pathway. TGFβ is considered a master regulator of epithelial to mesenchymal transition. To evaluate the association of TGFβ signaling with the upregulated gene expression profile and to TMZ/IR resistance, we utilized an inhibitor of the TGFβR1 serine threonine kinase. Animals bearing flank tumors derived from pre-treatment (P2) as well as recurrent tumors (R1) were randomized into four groups: 1) Control (untreated), 2) TMZ/IR treatment, 3) Treatment with TGFβ inhibitor alone (LY2109761), and 4) treatment with a combination of TMZ/IR and LY2109761. Western blot analysis of untreated tumor samples (P2 and R1) as well as treated recurrent tumor samples (RT1-5) revealed that treatment of tumors with LY2109761 resulted in decreased levels of THY1, ZEB2, CDK6, SOX2, and GLI2 compared to the untreated recurrent tumors (Fig. 4A). Furthermore, these markers of a mesenchymal/stem phenotype were also downregulated in tumors that received TMZ/IR and concurrent LY2109761. These findings demonstrate that LY2109761 appropriately targeted the pathways of interest as expected in our models of GBM.

**Fig. 4.**
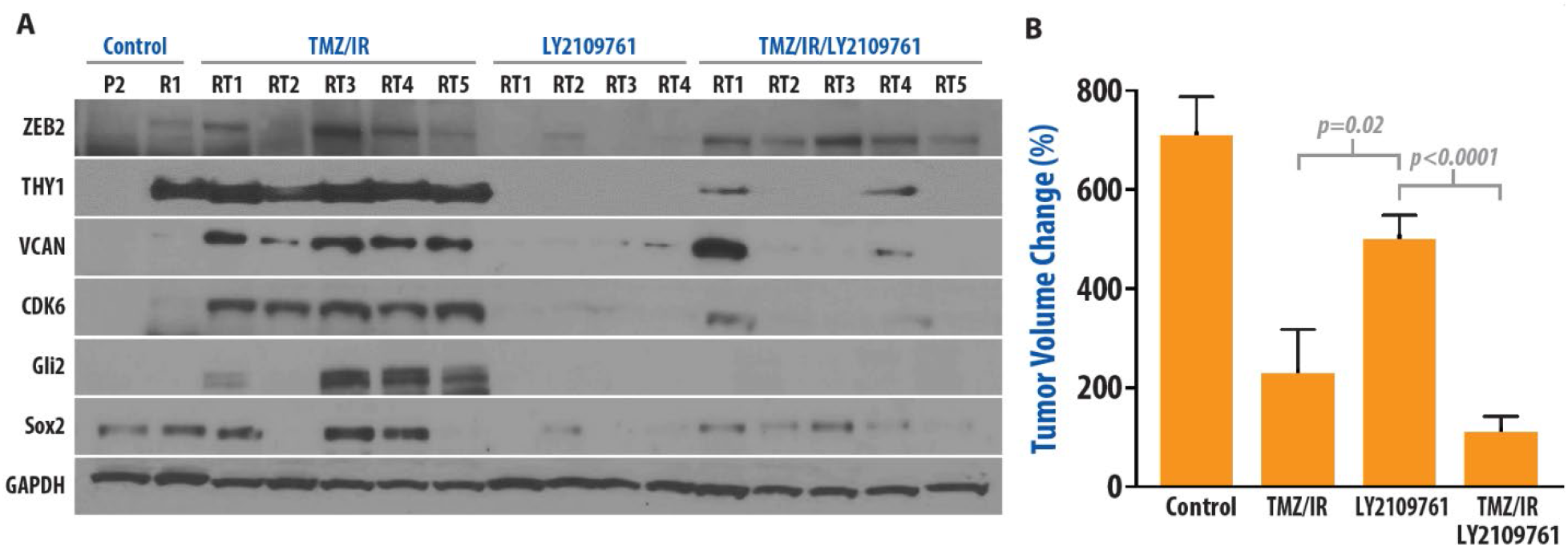
Inhibition of TGFβ signaling downregulates expression of mesenchymal and stem-like genes in recurrent tumors. **(A)** Western blot analysis shows that inhibition of TGFβ signaling results in a marked decrease in expression of proteins associated with a mesenchymal and stem cell signature in recurrent tumor samples. P2 (treatment naïve sample 2). R1 (recurrent sample 1), RT1-RT5 (recurrent samples treated as indicated in five independent animals) with TMZ/IR, LY2109761, or combination of TMZ/IR and LY2109761 as described below. **(B)** Animals bearing flank pre-treatment tumors as well as recurrent tumors were randomized into four groups 1)An untreated group (control), 2)Treatment with TMZ (TMZ, 5 days/week at 66mg/kg) and IR (5 days/ week at 2Gy) for two weeks followed by TMZ (3 days/week at 66mg/kg) every other week (TMZ/IR), 3)Treatment with LY2109761 where mice received 50mg/kg LY2109761 twice every day, 5 days/week for the first two weeks followed by 3 days/week every other week (LY2109761), and 4)TMZ/IR/LY2109761 group where the animals were treated with a combination of the three aforementioned treatment modalities. 5 replicates animals were evaluated. Mean volume + SEM shown.

Single agent treatment with LY2109761 did not significantly slow tumor growth (Fig 4B). However, combining LY2109761 with TMZ/IR substantially decreased tumor growth compared to single agent LY2109761 or TMZ/IR treatment (Fig. 4B). These findings suggest that TGFβ-mediated activation of mesenchymal and stem-like gene expression profiles promotes resistance to TMZ/IR, and that TGFβ inhibition downregulates the mesenchymal and stem-like phenotype restoring sensitivity to TMZ/IR.

### Rare THY1+ cell populations derived from treatment naive GBM are inherently TMZ/IR resistant, and repopulate the recurrent tumors

The above finding that *THY1* transcript and protein levels were upregulated within the recurrent tumor samples (more than 100-fold compared to treatment-naïve samples, Fig. 3A), combined with the observation that LY2109761 treatment resulted in decreased *THY1* expression and restored the sensitivity of recurrent tumors to TMZ/IR (Figs. 4A and 4B respectively), prompted us to investigate *THY1* further. In human cancers, *THY1* is associated with a poor prognosis, is linked with cancer stem cells and plays a role in adhesion, invasion, metastasis, immune evasion, and epithelial to mesenchymal transition[25–27]. Given the association of *THY1* expression with treatment resistance and a mesenchymal signature, its notable upregulation within the recurrent tumor samples, and its status as a cell surface marker, we quantitatively evaluated the presence of THY1+ cells in our tumors. In the treatment-naïve tumor biopsies we observed that THY1+ cells constituted 6-10% of the population. In contrast, 75-96% of the cells within recurrent tumors were THY1+ (Figs. 5A,B). FACS isolation and expansion of THY1+ cells from pre-treatment samples (Pre-THY1+), confirmed that they retained the expression of cell surface THY1(Figs. 5C,D). Pre-THY1+ cells were then implanted intracranially into nude mice and treated with TMZ/IR. These experiments revealed that the Pre-THY1+ cells were in fact inherently treatment resistant to a degree that mirrored the recurrent tumors (Fig. 5E). Additionally, the survival of mice bearing intracranial tumors derived using Pre-THY1+ cells was similar to mice having tumors derived from recurrent cells (Fig. 5F). This finding, combined with the consistent observation that each of the recurrent tumors (R1-R4, Fig 5A) exhibited an enrichment of THY1+ cells when compared to their respective pre-treatment biopsy (P1-P3, Fig. 5A) suggests that THY1+ cells represented a treatment recalcitrant cell population within the treatment-naïve tumors. These cells have stem cell like features with a mesenchymal phenotype, and likely survive therapeutic intervention and repopulate the tumor.

**Fig. 5.**
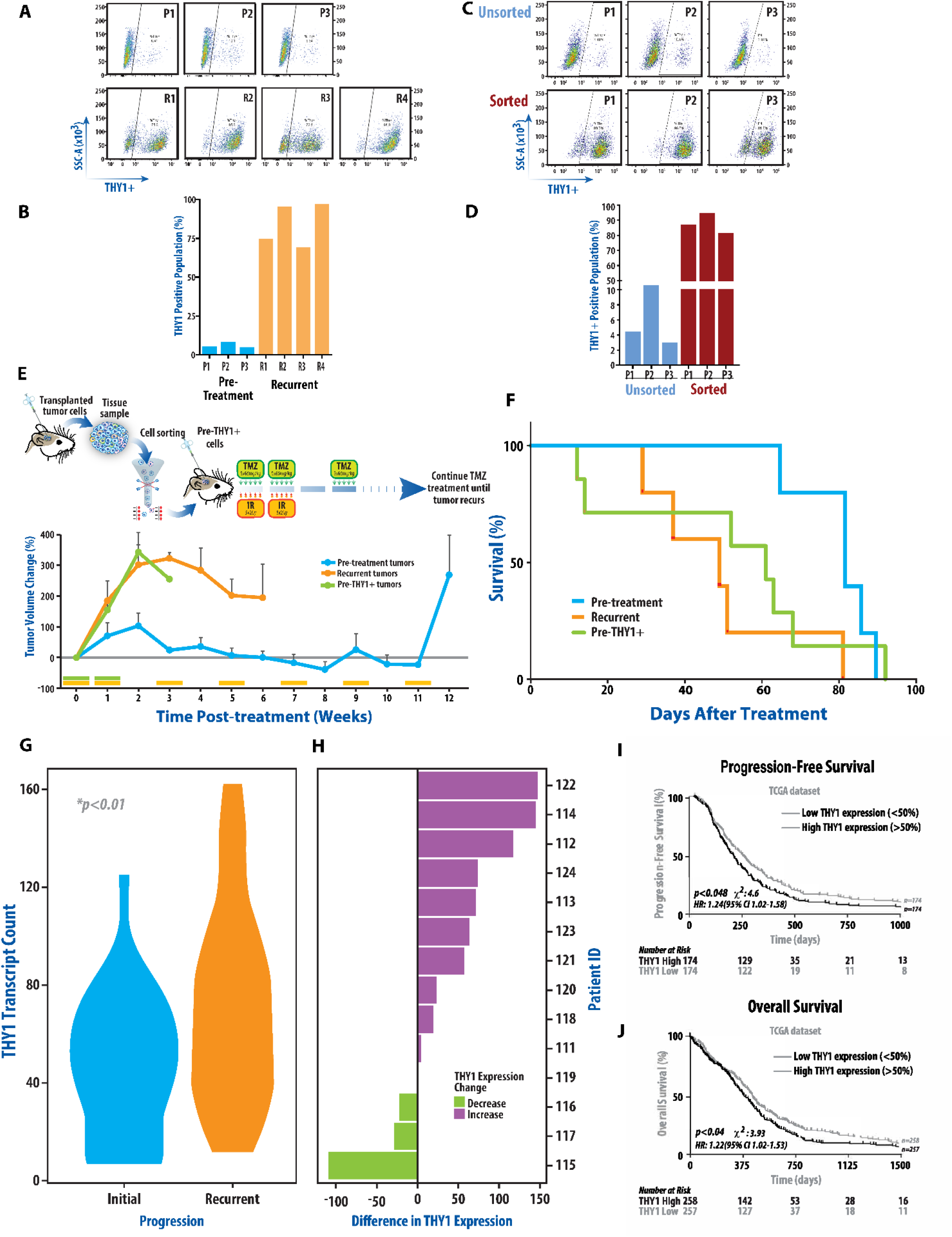
Rare THY1+ cell populations are identified in the pre-treatment samples, are inherently treatment resistant, and repopulate the recurrent tumors. **(A)** THY1 cell positivity is highly upregulated in recurrent samples as seen by flow cytometry and representative bar graph **(B)**. 6-10% of pre-treatment samples (P1-P3) demonstrate THY1-positivity while 72-96% of recurrent samples (R1-R4) reveal THY1-positivity(left) with quantification of replicate experiments shown(right). **(C, D)** THY1-positive cells were immuno-purified from P1, P2 and P3 samples by FACS. This sorted THY1+ cell population was cultured for expansion and re-tested to confirm stable THY1 cell surface staining. **(E)** Sorted THY1+ cell population from P2 (Pre-THY1+) was implanted intracranially to assess treatment response. Resistance to TMZ/IR was observed in the Pre-THY1+ tumor bearing animals that mirrored the phenotype observed in recurrent tumors, but distinct from pre-treatment samples. Mean tumor volume (**E**) (+SEM) and survival analysis by Kaplan-Meier (**F**) are shown. **(G)** Analysis of longitudinal GBM specimens reveals significant upregulation of *THY1* expression in recurrent tumors(GLASS consortium) with most patients showing increased expression **(H)**. Evaluation of TCGA clinical database reveals *THY1* expression is associated with significantly worse recurrence-free **(I)** and overall survival **(J)**.

To assess the clinical significance of upregulated *THY1* and its association with treatment resistance, we analyzed paired human GBM specimens from the GLASS consortium and other clinical databases which revealed that *THY1* gene expression is significantly upregulated in the majority of recurrent tumors (Figs. 5G,F; Fig. S4). Analysis of the TCGA database showed that high *THY1* expression was associated with significantly worse progression-free and overall survival (Figs. 5I,J). These findings illustrate that our model paralleled the clinical scenario, and further emphasize the role of *THY1* in the development of TMZ/IR resistance, and suggests that clonal expansion of THY1 expressing GBM cells likely contribute towards TMZ/IR resistance.

### *THY1* expression in human GBM is spatial niche-dependent

Our *in vivo* studies suggesting that *THY1* expression is associated with therapeutic resistance and a stem and mesenchymal gene expression signature, combined with recent studies suggesting that THY1+ cells often reside within a perivascular or stem cell niche and are associated with a mesenchymal and stem cell gene expression pattern [26, 28], prompted us to evaluate the spatial distribution of *THY1* expressing cells in patient-derived GBM tumor tissue. Spatial quantitative whole transcriptome analyses (spatial transcriptomics) of image-guided intraoperative human GBM biopsies from a region of high perfusion(i.e., with high vascularity) revealed that *THY1* expression significantly co-localized within the perivascular niche, characterized by co-expression of a glioma stem cell and mesenchymal signature, as well as TGFβ signaling (Fig. 6; Fig. S5A). We also performed spatial transcriptomics on a biopsy derived from a low perfusion region. This analysis also revealed that *THY1* was expressed within the perivascular niche, and was significantly co-localized with a mesenchymal and stem cell signature with upregulation of TGFβ signaling (Figs. 6D, F; Fig. S5B).

**Fig. 6.**
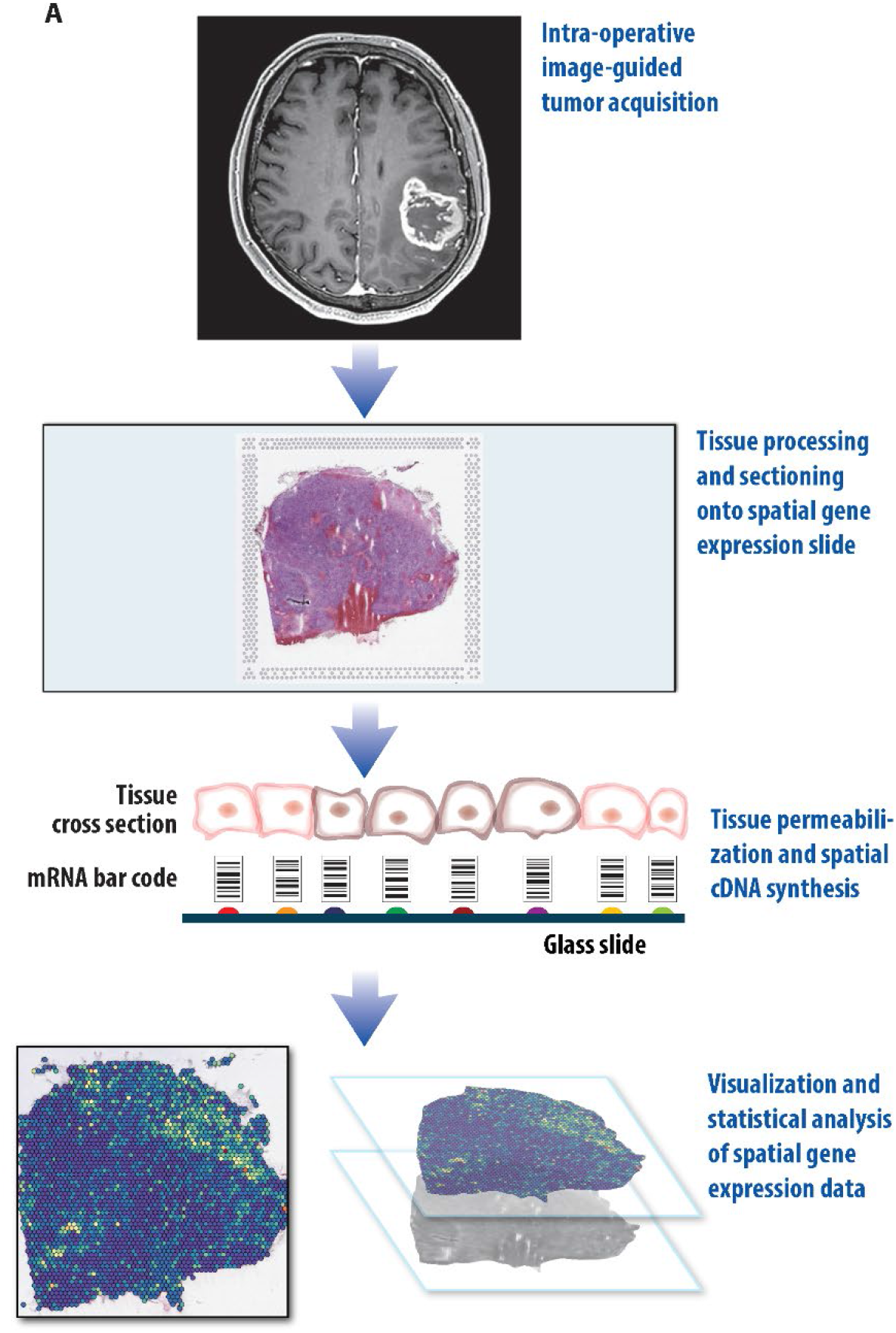
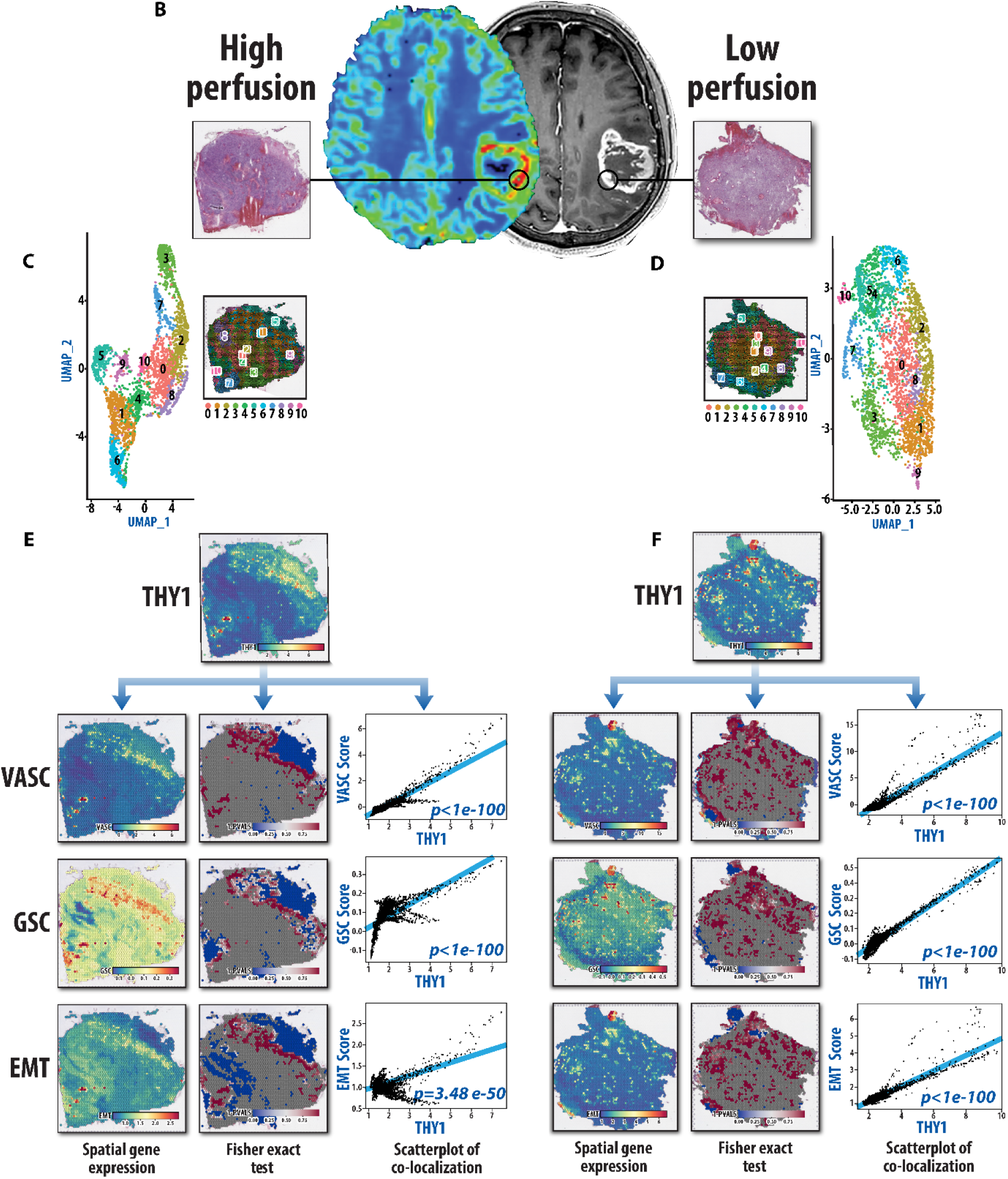
Spatial transcriptomic analysis (ST) of treatment-naive human GBM biopsies reveal that *THY1* expression significantly co-localizes with a glioma stem cell (GSC), and epithelial to mesenchymal transition (EMT) gene expression signatures within the perivascular niche. (**A**) A schematic depicting the experimental approach for evaluating the spatial distribution of gene expression profiles within a GBM biopsy. (**B**) MRI from a patient with primary GBM showing T1-post contrast weighted MRI and corresponding perfusion (CBV) scan. Two image-guided stereotactic biopsies were acquired, one from a contrast-enhancing region with relative high-perfusion, and a second from an enhancing region with low-perfusion. **(C,D)** UMAP analyses depicting spatial regions with unique gene expression profiles within both samples. **(E,F)** ST analysis of a GBM biopsy from a region with high perfusion **(E)** and Low Perfusion **(F)**. Spatial distribution of *THY1* expression is shown along with vascular (VASC), GSC, and EMT pathway enrichment scores. The left panels show the spatial gene expression of each individual gene set. The middle panel shows 1 minus p-values from a spatially local Fisher’s exact test within each spot for significant overlap between regions with high pathway enrichment scores and high THY1 expression. Significance in spatial overlap between the pathway enrichment scores and *THY1* expression was tested by using a sliding window approach to perform spatially-local Fisher’s test. On the right are scatterplots showing gene expression of *THY1*(x-axis) and enrichment scores for gene set of interest(y-axis) within each spatial location with line fitted. P-values show significance of correlation. *THY1* significantly co-localizes with these gene sets both in the high and low perfusion samples.

Since tumor-associated macrophages are known to drive malignant growth, stem cell maintenance, and epithelial to mesenchymal transition (EMT) [3, 29–32], we analyzed our spatial transcriptomic data which revealed that *THY1* also consistently co-localized with a macrophage gene expression profile (Fig. S5). We then performed gene ontology pathway enrichment analyses for THY1 high vs low regions which revealed that cell adhesion and extracellular matrix pathways are highly upregulated in THY1-enriched regions both in the hypervascular(Fig. S6A,C) and hypovascular(Figs. S6B, D) samples, supporting our *in vivo* analyses (Fig. 2C). These data show that THY1+ cells consistently localize within a common niche, suggesting a critical role within the tumor microenvironment in a signaling axis that contributes to maintenance of a mesenchymal and cancer stem cell phenotype which contributes to treatment resistance.

## DISCUSSION

Despite aggressive therapy for GBM with maximal safe resection and TMZ/IR, recurrence is inevitable, and therapeutically actionable mechanisms that drive resistance have yet to be identified. To characterize the molecular basis for the development of therapeutic resistance in GBM, we have developed and analyzed a pre-clinical *in vivo* recurrence model along with spatial analyses of patient-derived biopsies. A key finding from the described studies indicates that a rare population of pre-existing, therapy-resistant tumor cells within treatment-naïve GBM likely drive tumor recurrence. Specifically, this THY1+ cell population resides within a perivascular tumor niche strongly expressing EMT and stem cell signatures, and co-localizes with macrophages. Our findings also suggest that tumor recurrence may be driven through critical cell-cell interactions, between THY1 expressing cells and macrophages, within the perivascular tumor microenvironment that drive a mesenchymal transition.

The GBM recurrence model that we have developed recapitulates the clinical treatment paradigm and results in profoundly resistant tumors at recurrence. Most notable was the finding that THY1 expression was increased greater than 100-fold in recurrent biopsies compared to pre-treatment biopsies. We then isolated, using THY1 as a cell surface marker, a population of cells from treatment-naïve tumors that recapitulated the treatment resistance phenotype of the recurrent tumor samples. The biological significance and clinical relevance of these results is further emphasized by the finding that THY1 expression is significantly upregulated in the majority of patients at the time of recurrence, and that increased THY1 expression was associated with worse progression-free and overall survival. Although clonal selection of pre-existing therapy-resistant cells in GBM has been described previously[8, 33–35], the findings presented here provide a strong proof of the concept and identifies a specific phenotype and gene expression profile that characterizes this cell population.

Gene expression analysis of replicate paired pre-treatment and recurrent tumors from the same mouse revealed that development of a resistant phenotype was also associated with elevated expression of genes related to mesenchymal transition, stemness, cell adhesion and components of the extracellular matrix, as well as downregulation of genes associated with cell death. A mesenchymal gene expression signature results from the biological process of epithelial– mesenchymal transition (EMT) and is characterized by an enhanced capacity for invasion and metastasis in many malignancies [6, 36–38]. Here we nominate THY1 as among the most significantly upregulated membrane proteins in cells acquiring a mesenchymal phenotype with stem cell properties in GBM [24-27, 39, 40]. THY1 is a cell surface receptor that has been identified in several nonpathological cell types and is critical for cell-cell communication as well as cell-matrix interactions resulting in many physiological processes including proliferation, adhesion, and migration[24, 39–41]. THY1 has been identified in several malignancies including renal cell cancers, breast cancer, hepatocellular carcinomas, ovarian cancer, and pancreatic tumors[28, 40–45] and has been associated with invasion, metastasis, immune evasion, and poor prognosis, all of which are hallmarks of EMT.

In our paired *in vivo* analyses, the recurrent samples which had THY1 upregulation also had significantly increased expression of genes associated with cell morphogenesis, cell-cell contact, as well as cell adhesion and extracellular structure organization. These results were corroborated by our gene ontology pathway enrichment analyses of human GBM specimens (Fig. S6). These cellular pathways mediate signals for growth, migration, survival, differentiation, and resistance to cell death [46, 47]. Avril *et al* demonstrated that THY1/CD90+ tumors have increased association with invasiveness and multi-focal tumor presentation[41]. These studies further supports a role for THY1 in cell-cell communication and migration mediating treatment resistance.

Therapeutic resistance in glioblastoma is driven by marked inter- and intra-tumoral heterogeneity both at the cellular and microenvironmental level[3, 7, 48–50]. In the context of the tumor microenvironment (TME), distinct tumor niches, especially perivascular, hypoxic, and invasive tumor niches, have been shown to be enriched with therapy-resistant cancer stem cell populations [3, 9, 10, 49–53]. These niches support resistant populations and play a critical role in tumor recurrence, as the interplay between the treatment refractory cells and adjacent cells including immune, stromal, differentiated cancer, glial, neuronal, and endothelial cells drives quiescence, invasion, immune suppression as well as other features promoting treatment resistance[3, 54–56]. In order to target these stem-like, therapy-resistant cells, the signaling cascades that promote a resistant phenotype in the context of the local TME need to be better understood. The recently developed technique of spatial transcriptomics (ST)[57, 58] provides the ability to interrogate whole transcriptome expression analysis *in situ*, permitting the assessment of the local tumor microenvironment. Due to the robust and reproducible evidence that THY1-positivity serves as a marker for a therapy resistant phenotype and hence may serve as a spatial landmark, we performed ST analyses of image-guided biopsies of a patient-derived GBM. This analysis was performed in an unsupervised and spatially-naïve manner and gene expression data was then superimposed onto the original H+E imaging. We found that in the analyses of hyper-perfused and hypo-perfused regions of a single GBM, THY1 expression consistently co-localized with a mesenchymal and stem-like signature and was found primarily in the perivascular niche [26], validating our studies in mouse models. Furthermore, our ST studies revealed co-localization of THY1 expression with a macrophage gene expression signature. Tumor-associated macrophages (TAMs) have been shown to maintain stem cell populations[3, 29, 48, 59–62], and promote invasiveness and immune evasion through numerous interactions. For example, in breast cancer, TAMs were shown to preferentially infiltrate THY1+ regions, bind the THY1 receptor on cancer stem cells which resulted in juxtracrine signaling promoting invasiveness and supporting a stem cell niche, with activation of SRC and NFκB signaling cascades[28]. These studies further revealed that depletion of macrophages resulted in arrest of tumor growth. In glioma, a recent study demonstrated that increased THY1 expression correlated with SRC phosphorylation, and resulted in a SRC-dependent tumor cell migration as well as cell-matrix interaction which was inhibited by the SRC-inhibitor, dasatinib[41]. Interestingly, a recent study found that local THY1+ mesenchymal stem cells induced glioma progression through increasing tumor proliferation and cell migration[27]. Our data, in conjunction with existing literature, highlight the importance of cell-cell interactions within the TME in tumor progression and the critical role of THY1 in paracrine/juxtracrine signaling within its local niche.

Our *in vivo* recurrent tumors displayed a mesenchymal gene expression profile characteristic of TGFβ pathway activation, which was further validated by ST analyses of our patient-derived tissues. A number of studies have independently demonstrated a role for TGFβ signaling in treatment resistance [63–67], expression of stem cell markers, neurosphere formation ability as well as tumorigenic potential in mouse models [66, 68, 69]. Furthermore, inhibition of TGFβ signaling has been associated with a depletion of the stem cell population in breast cancer, leukemia and glioblastoma[68–72]. Our findings support this as inhibition of TGFβ signaling using LY2109761 decreased the expression of mesenchymal and stem-like genes. Furthermore, treatment of recurrent tumors with LY2109761 restored sensitivity to TMZ/IR. Importantly, although LY2109761 had minimal efficacy as single agent, we observed a significant restoration in sensitivity to standard of care TMZ/IR when the two modalities were combined. This finding directly implicates the TGFβ signaling pathway in imparting a mesenchymal, stem-like, therapy-resistant phenotype as previously described[72]. Recent clinical trials investigating various TGFβ inhibitors have not shown promise but have typically utilized TGFβ inhibitors alone or in combination with other drugs including lomustine [73, 74]. However, our findings suggest that TGFβ inhibition may be most effective in a recurrent setting to reverse the resistance to TMZ/IR, especially if patients are enriched for the expression of a mesenchymal and stem-like signature described here. The specific pathway to resistance likely only occurs in a subpopulation of patients as evidenced by Wang *et al* who found mutations in LTBP4, a known activator of the TGFβ pathway, in 11% of patients at recurrence [8]. Therefore, identifying subgroups of patients who are likely to benefit from TGFβ targeting is critical[65]. Activation of the TGFβ signaling pathway and a mesenchymal phenotype is also associated with enhanced capacity to repair DNA damage thereby leading to chemo- and radiation-resistance through several transcription factors [75]. For example, ZEB2, which was upregulated in our recurrent samples, has been shown to affect ATM/ATR activity upon DNA damage [76]. Its homolog ZEB1, promotes radioresistance in breast cancer by regulating the stability of CHK1 and regulates TMZ sensitivity via regulation of MGMT in glioblastoma [77, 78].

The development of recurrence is a dynamic process that is poorly understood. Many have theorized that the evolutionary pressure during treatment results in selection of clonal cells versus expansion and mutation of a subclonal population of cells [5, 33, 34, 79]. Our findings demonstrate that evolution and growth of a pre-existing tumor-cell sub-clonal population can also lead to recurrence. Chowell *et al*, demonstrated that treatment-naïve tumors often harbor rare treatment-resistant subclones which are undetectable with standard assays[33]. A study by Kim *et al* found that the genetic path to tumor recurrence can be unique and difficult to predict when examined across multiple samples [5]. Others have demonstrated that various subclones from a single tumor can exhibit variable treatment responses *in vitro*, and asserted that this information can be used to predict clinical response[80].

Finally, our study provides a distinct clinically relevant mouse model to study treatment resistance in GBM and defines a rare cell population, expressing THY1 that is intrinsically resistant to TMZ/IR therapy. This approach provides a unique opportunity to explore mechanisms for tumor resistance and therapeutic interventions critical for patient care. Although *in vivo* immunocompromised murine models do not perfectly replicate the clinical paradigm or the human tumor microenvironment, our findings are validated by clinical findings. Furthermore, our human spatial transcriptomic analyses provide additional insights into interactions between human tumor cells and the tumor microenvironment. The use of patient-derived explants rather than immortalized cell lines in an intracranial mouse model to evaluate the evolution of resistance to combination therapy using a genome wide analysis of gene expression also provides confidence that our findings approximate the clinical setting. Consistent with clinical experience, our results demonstrate that adjuvant TMZ prolonged the overall survival from 7 weeks to 13 weeks. The development of recurrence in animals while undergoing adjuvant TMZ suggests that the tumors were truly resistant. In addition, the demonstration that resistance to TMZ/IR was recapitulated upon re-implantation of recurrent tumor tissues confirms this conclusion. Further efforts are needed to define how inherently resistant cells drive recurrence through interactions within the perivascular niche, which may lead to the identification of pathways to target for clinical benefit. In the recurrence model described here, we have utilized a combination of TMZ/IR to evaluate the mechanisms of treatment resistance. Although combination TMZ and IR is standard of care for GBM patients, a number of reports have tested the molecular basis for GBM resistance utilizing a single modality only [81, 82], which although important, does not mimic the emergence of resistance in clinical settings.

## Acknowledgments

Tumor specimens obtained from the University of Michigan were obtained through the CNS Tissue Biorepository (HUM00024610) and the Henry Ford brain tumor center. The results shown here are in part based upon data generated by the TCGA Research Network: https://www.cancer.gov/tcga, and the Glass Consortium (https://www.glass-consortium.org/). We would like to thank Steven Kronenberg for assistance with editing and illustration of figures. We would also like to thank Kait Verbal for her assistance in acquiring tumor samples through the CNS Tissue Biorepository. We would like to thank Dah-Luen Huang, Pavlina Zafirovska, Susan Dagenais, Judy Opp, Ingrid Bergin, and Olivia Koues for assistance in processing our ST samples.

## Author contributions

Conceptualization: W.N. Al-Holou., H. Wang, A.C. de Carvalho, T.Mikkelsen, A.Rehemtulla

Methodology: W.N. Al-Holou., H. Wang, V. Ravikumar, M. Oneka, R.G.W. Verhaak, A. Rao, H. Kim, A. Rehemtulla

Investigation: W.N. Al-Holou., H. Wang, V. Ravikumar, M. Oneka, R.G.W. Verhaak, H. Kim, D. Pratt, S. Camelo-Piragua, A. Rao, C. Speers, A. Rehemtulla

Funding acquisition: W.N. Al-Holou, A.C. de Carvalho, T. Mikkelsen, A. Rao, A.Rehemtulla

Tissue and/or cell contribution: W.N. Al-Holou, A.C. de Carvalho, T. Mikkelsen, T.Hollon, O. Sagher, J.A. Heth, K.M. Muraszko

Writing – original draft: W.N. Al-Holou., H. Wang, V. Ravikumar, A. Rao, M. Oneka, A. Rehemtulla

Writing – review & editing: W.N. Al-Holou., H. Wang, V. Ravikumar, M. Oneka, R.G.W. Verhaak, D. Pratt, S. Camelo-Piragua, C. Speers, D.R. Wahl, S. Shankar, T.Hollon, O. Sagher, J.A. Heth, K.M. Muraszko, T.S. Lawrence, A.C. de Carvalho, T.Mikkelsen, A. Rao, A. Rehemtulla

## Data Availability Statement

The data generated in this study are available upon request from the corresponding author.

---

**Fig. S5.**
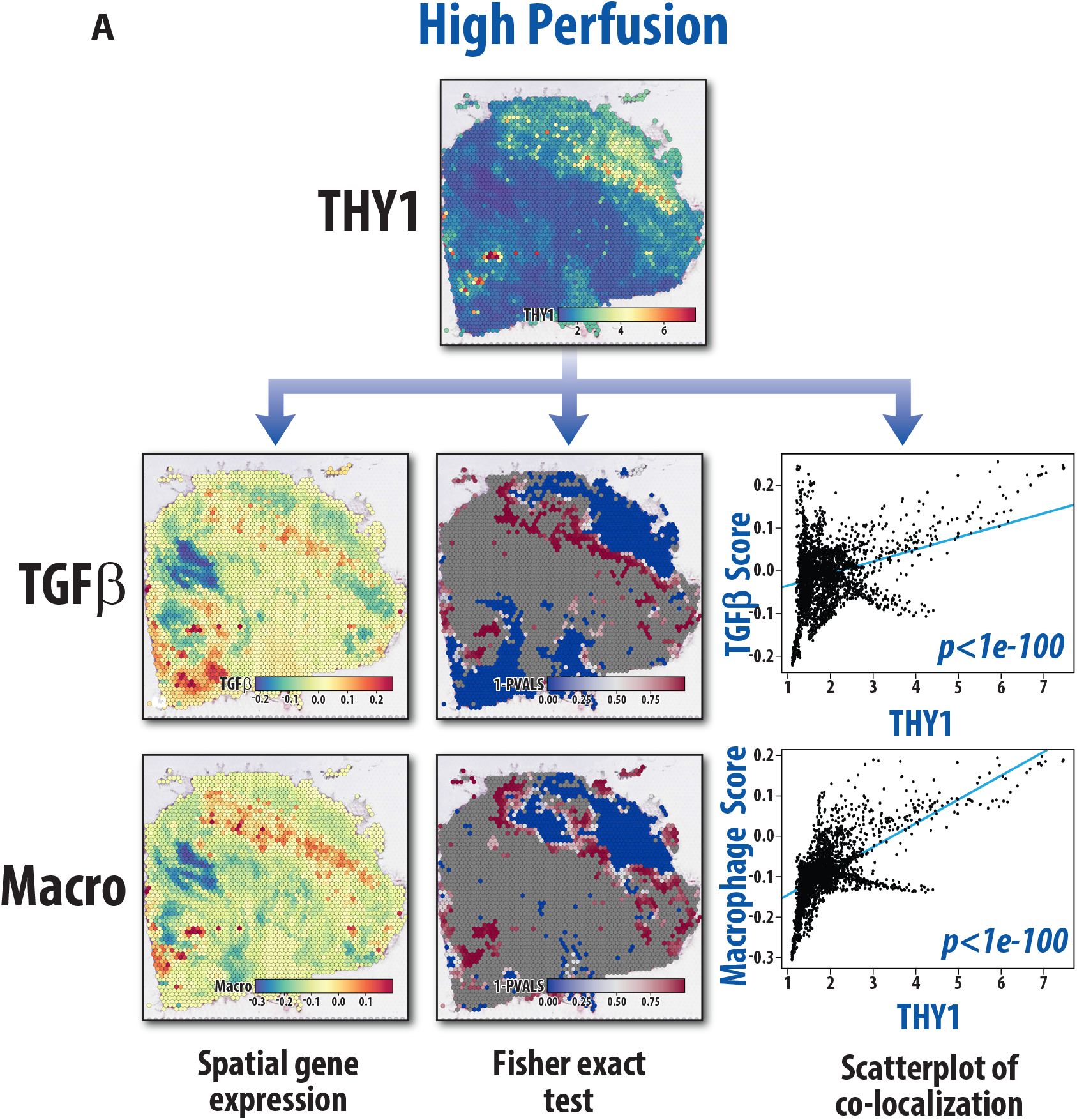
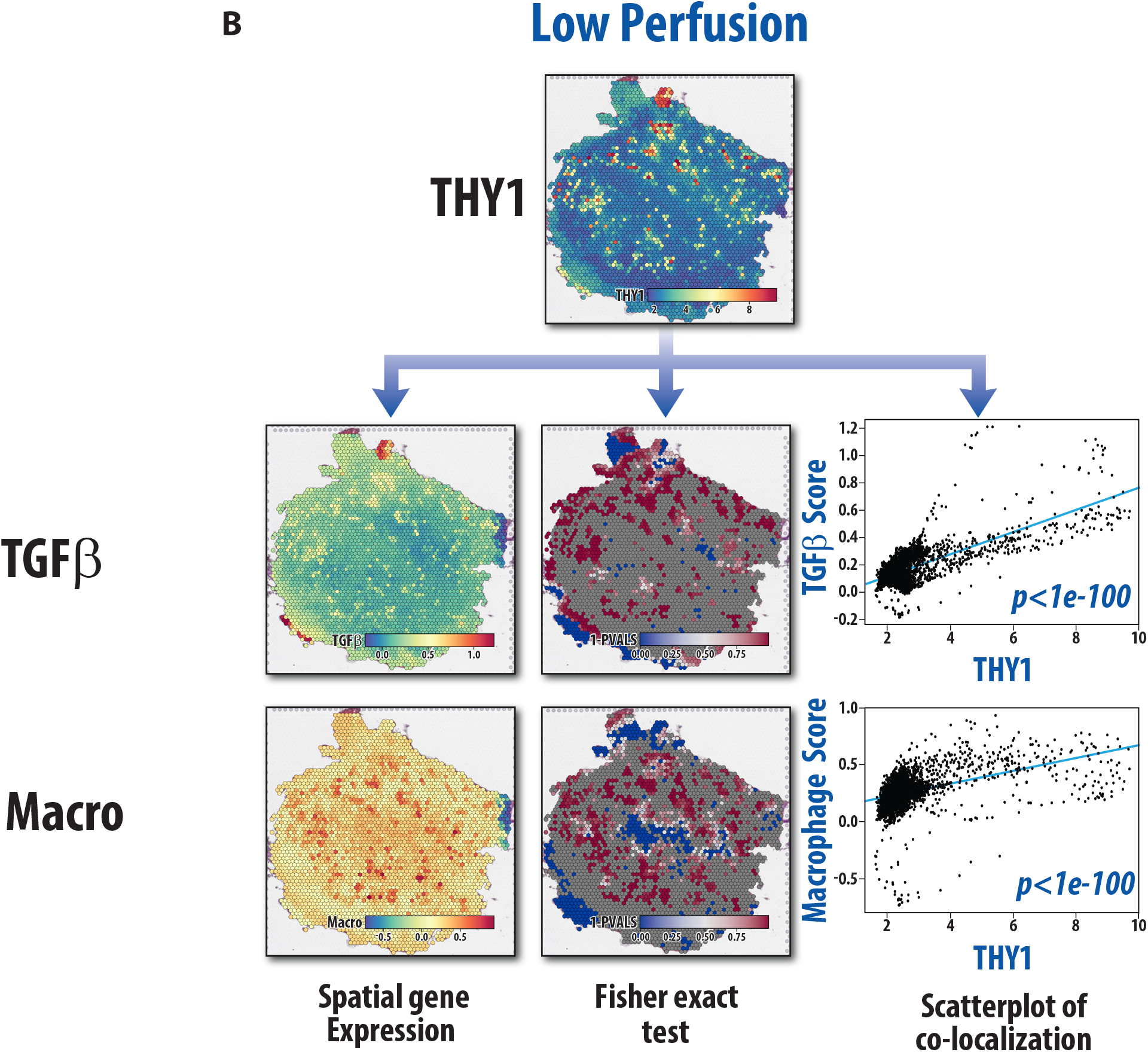
Spatial transcriptional (ST) analyses of human GBM reveal that *THY1* significantly co-localizes with increased TGFβ pathway and macrophage gene expression signatures. MR-image guided biopsies from high perfusion **(A)** and low Perfusion (**B**) regions of a human GBM shows that *THY1* expression co-localizes with TGFβ signaling and macrophage gene enrichment signatures. On the left is shown the spatial gene expression of each individual gene set. In the middle are spatial Fisher exact test showing p-values for each region comparing THY1 expression along with gene sets of interest. Significance in spatial overlap between the pathway enrichment scores and THY1 expression was tested by using a sliding window approach to perform spatially-local Fisher’s test. On the right are scatterplots showing gene expression of *THY1*(x-axis) and gene set of interest(y-axis) within each spatial location with line fitted. P-value shown. These results validate our *in vivo* findings showing TGFβ signaling pathway upregulation in THY1+ samples, and support existing literature suggesting THY1+ tumor cells interact with tumor infiltrating macrophages(*20*).

